# Metabolic model predictions enable targeted microbiome manipulation through precision prebiotics

**DOI:** 10.1101/2023.02.17.528811

**Authors:** Georgios Marinos, Inga K. Hamerich, Reena Debray, Nancy Obeng, Carola Petersen, Jan Taubenheim, Johannes Zimmermann, Dana Blackburn, Buck S. Samuel, Katja Dierking, Andre Franke, Matthias Laudes, Silvio Waschina, Hinrich Schulenburg, Christoph Kaleta

## Abstract

The microbiome is increasingly receiving attention as an important modulator of host health and disease. However, while numerous mechanisms through which the microbiome influences its host have been identified, there is still a lack of approaches that allow to specifically modulate the abundance of individual microbes or microbial functions of interest. Moreover, current approaches for microbiome manipulation such as fecal transfers often entail a non-specific transfer of entire microbial communities with potentially unwanted side effects. To overcome this limitation, we here propose the concept of precision prebiotics that specifically modulate the abundance of a microbiome member species of interest. In a first step, we show that defining precision prebiotics by compounds that are only taken up by the target species but no other species in a community is usually not possible due to overlapping metabolic niches. Subsequently, we present a metabolic modeling network framework that allows us to define precision prebiotics for a two-member *C. elegans* microbiome model community comprising the immune-protective *Pseudomonas lurida* MYb11 and the persistent colonizer *Ochrobactrum vermis* MYb71. Thus, we predicted compounds that specifically boost the abundance of the host-beneficial MYb11, four of which were experimentally validated *in vitro* (L-serine, L-threonine, D-mannitol, and γ-aminobutyric acid). L-serine was further assessed *in vivo*, leading to an increase in MYb11 abundance also in the worm host. Overall, our findings demonstrate that constraint-based metabolic modeling is an effective tool for the design of precision prebiotics as an important cornerstone for future microbiome-targeted therapies.

## 3. Introduction

While traditionally considered commensals, it is becoming increasingly clear that the microbial communities living within and on higher host organisms (i.e., the microbiome) are integral parts of host physiology. These microbial communities fulfill essential roles in pathogen protection ^1^, immune system education ^2^, production of vitamins ^3^, metabolism of xenobiotics ^4,5^ and increase the plasticity of their hosts to respond to evolutionary challenges ^6,7^. Thus, the composition of the microbiota plays an important role in the well-being of hosts and its ability to respond to environmental challenges ^8,9^, as indicated by the frequent shift of microbiota composition observed in the context of diseases ^10^ and exposure to environmental stressors ^11^. These observations and work in animal models have led to the development of the notion that the composition of the microbiome itself might be an important modulator of host health ^12^ as well as fitness and that a targeted modulation of microbiome composition might offer a completely new avenue for modulating the host.

It has been shown that dietary patterns (e.g., western diet) can affect the structure of the microbiome, however their effect is transient ^13^. Besides an influence of dietary uptake on microbiome composition, various approaches to modify the microbial communities of organisms have been used ^14^: the transfer of entire microbial communities from a donor to a recipient host (fecal transfer), the provision of compounds used by particular microbial species (prebiotics), the provision of microbial strains of natural or synthetic origin (probiotics), the provision of compounds of microbial origin (postbiotics) and combinations thereof (synbiotics). While fecal transfer is the most commonly used approach to modulate microbial community composition in animal models, it involves the transfer of entire microbial communities and therefore not only desirable traits. In consequence, there is only a single medical indication in which fecal transfers are routinely used thus far ^15^ and there are potential concerns about the safety and stability of the engraftment of transplants ^16^.

Although the concept of supplementation with pre-, pro- or synbiotics has been commonly proposed as a method of microbial manipulation, these approaches are still severely limited. In humans, oligosaccharides are proposed as prebiotics to alter the microbial community on both the structural and functional level, since they can be fermented by microbial species, which in turn can proliferate and produce bioactive compounds such as short-chain fatty acids ^17^. However, there is yet only little knowledge about other classes of compounds that might alter the microbial composition. Another important aspect to consider is the specificity with which individual target microorganisms can be manipulated using prebiotics. Thus, while oligosaccharides are considered beneficial there is often uncertainty about which species can degrade them, as multiple, unrelated species can degrade similar types of oligosaccharides (e.g., *Bacteroidetes, Lactobacilli*, and *Bifidobacterium* species can degrade fructans) ^17,18^. In case of supplementation with probiotics and synbiotics, other concerns have been raised. For instance, the ability of specific Bifidobacteria to stably colonize the human gut was only reported for ⅓ of the participants in a clinical trial ^19^. Moreover, introducing exogenous bacteria can have unwanted side effects. For example, some of the *Bacillus* species which were used as probiotics in aquacultures were found to carry virulence and antibiotic resistance genes ^20^. Hence, to overcome these limitations in current microbiome manipulation approaches, we need to focus on prebiotics with well-controlled modes of action (e.g., microbial pathways of prebiotics degradation) that preferentially modulate the abundance of resident microbial species. Since mechanisms of pathogenicity are often complex and context specific ^21^, it is preferred if these species are native to the microbiome of the host, as unwanted side-effects due to the introduction of foreign bacterial species might be avoided. Within this work, we will refer to prebiotics that specifically modulate the abundance of selected bacterial strains within the microbiome of a specific host as “precision prebiotics”.

To design and evaluate such precision prebiotics, it is essential to start with a tractable model system such as the well-established model organism *Caenorhabditis elegans*. The native microbial community of the invertebrate *C. elegans* was first described in 2016 ^22^, yet the nematode has already been used for over twenty years as a model for studying host-microbial interactions ^23^. To similarly promote *C. elegans*-microbiome research, a synthetic 12-member *C. elegans*-specific microbiome resource, *CeMbio*, was recently established ^22^. Part of this consortium is the Gram-negative *Pseudomonas lurida* MYb11 that is known to possess antifungal ^24^ and antimicrobial properties, thereby providing protection against the pathogen *Bacillus thuringiensis* (Bt) ^25,21^. Besides its protective effects, MYb11 is known to colonize and persist in the *C. elegans* intestine ^22^ and thus increase the nematode’s fitness during infection ^25^. *Ochrobactrum vermis* MYb71 is another member of the CeMbio community ^22^ and affects different host life-history traits ^22,26^, persistently colonizes the nematode ^24^, and has been described as competitive against other bacteria in terms of space and nutrients ^27^. Both bacteria are also able to interact metabolically with the nutrient environment determining their type of interaction ^27^. Hence, a microbial community consisting of one beneficial bacterium and another one that is competitive offers an excellent opportunity for developing targeted approaches for microbiome manipulation.

A key prerequisite for the development of precision prebiotics is a detailed understanding of the metabolic functions present within a microbial community and how specific metabolic interventions affect microbiome metabolism. One important approach to investigate metabolic functions in microbial communities are constraint-based metabolic modeling approaches ^28–30^. Based on the genetic information either on the level of each bacterium or from the overall community and biochemical databases, these methods allow one to simulate the network of chemical reactions that represent the metabolic reactions of the microorganism(s) or the community ^31^. To this end, genome-scale metabolic models of the bacterial species within a microbiome have to be reconstructed from genomic data. These models summarize the metabolic functions present within an organism and can be reconstructed using automated reconstruction approaches ^32,33^ or obtained from existing model repositories ^34^. Furthermore, accounting for the growth environment of a species ^35^, optimization approaches such as flux balance analysis (FBA) for individual species ^28^ or constraint-based community modeling approaches ^36,37^ can be used to predict metabolic activities within individual species as well as interactions between species. These methods have already been used to identify metabolic alterations in the microbiome of Parkinson’s disease patients ^38^ as well as patients with inflammatory bowels disease ^39,27^ and to study the interaction of *C. elegans* as well as its microbiota in the context of drug-microbiome-host interactions ^30^. For *C. elegans*, metabolic models of the *CeMbio* community including MYb11 and MYb71 ^22^ and other members of the microbiota are available ^27^.

In this work, we used metabolic networks of MYb11 and MYb71 also incorporating phenotypic information from *Biolog* growth assays in a computational screen for precision prebiotics that are able to specifically boost the abundance of the host-protective species MYb11 over MYb71. We tested four of the predicted compounds, L-serine, L-threonine, D-mannitol, and γ-aminobutyric acid (GABA) *in vitro* and confirmed that they selectively boost the abundance of MYb11. Moving to the *C. elegans* host system, we moreover showed that L-serine is able to selectively boost the abundance of MYb11 when the *C. elegans* host was colonized with both bacterial species. Thus, we demonstrate that constraint-based microbial community modeling approaches could represent a key tool for the design of targeted microbiome intervention approaches.

## 4. Material and Methods

### a. Computational Methods

#### Identification of unique uptake compounds

We explored to what extent individual metabolites, which are only taken up by a single bacterial species and no other, could serve as potential supplements to target individual species. To this end, we considered gut microbiome data from a Kiel-based human cohort comprising 1280 participants for which 16S rRNA gene sequencing data is available, which we had utilized previously ^30^. As reported previously, the reads were mapped to the 16S sequences of the genomes of bacterial species present in the AGORA bacterial collection ^34^, the read counts of each sample were normalized to a sum of 1 and removal of species below 0.1% abundance in each sample. We used *gapseq* version 1.1 ^32^ on the genomes of the identified species assuming the averaged dietary uptake of the human cohort ^30^ as input to the models to reconstruct genome-scale metabolic models. Subsequently, we identified for each bacterial species which metabolites can be taken up. To this end, we minimized the flux through each exchange reaction using flux balance analysis (FBA) ^40^ implemented in the R-package sybil ^41^ which corresponds to maximizing the uptake of that metabolite. If the minimal flux through the exchange reaction was non-zero this metabolite could be taken up. Having identified those compounds for each species, we determined for each individual bacterial species in the gut microbiome of each participant whether it could take up a compound that could not be taken up by any other species in the bacterial community of this participant. Finally, we determined for each participant which fraction of the bacterial species contained in the gut microbiome had at least one unique uptake compound. Similarly, we reconstructed 78 bacterial models belonging to the *C. elegans* microbiota based on their published genomes ^27^ using *gapseq* version 1.2 ^32^. We then created random subsets of the models with increasing size (from 2 to 78, 50 iterations for each size) and determined the portion of strains with unique uptake compounds for each community. For the CeMbio community we used metabolic models reconstructed with gapseq 1.2 ^32^ from Zimmermann *et al*. (in preparation).

#### Computational supplementation experiments

In our simulations, we utilized metabolic networks of *Pseudomonas lurida* MYb11 and *Ochrobactrum vermis* MYb71 that were created from their genome sequences ^27^ using *gapseq* ^32^ version 1.1. Information from phenotypic microarrays about the organisms’ metabolic capacities was integrated into the models using the ‘adapt’ module of *gapseq*. Specifically, we incorporated data from the *Biolog* EcoPlate (assuming growth if OD590 - OD750 > 0.1) ^22^. During growth simulations, exchange fluxes of the models were constrained by assuming a computational Nematode Growth Medium (NGM) ^30^ and aerobic conditions. These constraints can be found in Supplementary File (Sheet: S1). Additionally, the inflow of copper and iron cations was increased, so that they did not restrict the growth of the models.

We used three distinct approaches for modeling individual microbes and microbial communities. To model the growth of each bacterium independently, we used FBA ^40^. We also utilized individual-based modeling based on *BacArena* ^36^ R-package and a community-FBA as described previously ^39,42,43^. The community-FBA approach is implemented in the R-package *MicrobiomeGS2* (www.github.com/Waschina/MicrobiomeGS2). In all types of simulations, we focused on the identification of nutritional supplements that boosted the growth of the target species MYb11.

For the individual supplementation experiments, the metabolic models of both organisms were constrained with the NGM computational diet, assuming that the molecular values of the dietary compounds (molecular quantities) can be directly incorporated into the models (as fluxes in the lower bounds of the models) without any transformation. This is based on the assumption that the uptake rate of compounds is depicted by its maximum available amount ^36^. Supplementations for each metabolite were conducted by increasing the possible influx of the supplemented metabolite by 10 mmol/gDW/hr. For testing the effects of supplements on single growth, the relative increase in growth rate compared to growth without supplement for each model was recorded. To prevent any numerical instabilities, all raw values were rounded to the sixth digit and relative changes bigger than 0.01 were taken into account.

For simulations using community FBA, the metabolic models of both organisms were merged into one microbial community model. Please note that we did not use a fixed community biomass composition as previously ^30^, since we employed community FBA to predict the resulting growth rates of individual species after maximizing the growth of the community. Thus, the community model was constrained with the NGM diet - extended with the individual supplements, as described in the previous paragraph. The growth rates of individual species were predicted by maximizing the sum of the growth rates of individual species with concomitant minimization of the sum of total fluxes as implemented in the R-package sybil ^41^ (with a coefficient of -10^−6^ for the sum of fluxes in the objective function). Since the supplementations usually also increased the growth of the entire community, the predicted growth rates of each species were normalized with the respective growth rate of the community. Predicted growth rates following supplementation were contrasted against growth rates without supplementation to identify compounds whose effect was larger for the growth of MYb11 in comparison to MYb71. To exclude any numerical artifacts, rounding to the sixth digit and a cut-off of 0.01 in compound selection were also applied in this section.

For individual-based modeling of microbial communities, the metabolic models of both species were simulated in a virtual environment in *BacArena*, where they could randomly move and reproduce ^36^ on a 30 × 30 grid. As a growth medium we used the NGM diet (in mM) which we diluted 1000-fold, because otherwise bacterial models overgrew the entire simulation environment before nutrients were exhausted. For the simulations, we inoculated the growth environment with 20 individuals of each species and simulated bacterial growth for 12 iterations with 15 replicates for each supplementation using FBA. A secondary objective to minimize the total flux while maximizing growth rates of individuals was applied. To simulate the supplementation experiments, 0.01 mM of each compound of interest was added once in the beginning of the simulation. To identify significant changes in the growth profiles of individual species following each supplementation, we extracted the cell biomass information from the simulations using the command *plotGrowthCurve* and we calculated the relative change of the total biomass values (time step 13; rounding to the sixth digit) due the supplementations for each model. Then, the relative changes between the bacterial models were compared using Wilcoxon rank sum tests ^44^. The resulting p-values were corrected for multiple testing using false discovery rate control implemented in R. We selected those compounds whose effect on MYb11 was larger than on MYb71 based on the median of the respective relative changes for each model.

Based on these simulation techniques, we identified those compounds that specifically boosted MYb11 across the simulations. Subsequently, we subsetted this list to compounds for which we had positive confirmation of their uptake by MYb11 from Biolog experiments ^22,27^. For the enrichment analysis of growth-supporting compounds of MYb11, we used the human metabolome database (HMDB) annotation for the available metabolite classes ^45^. Briefly, we tested whether the sets of candidate compounds were enriched in the metabolite classes defined in HMDB using Fisher’s exact test. As the underlying set of metabolites, we used the metabolites contained in the metabolic model of MYb11 in the different metabolite classes.

#### System information and software

Simulations and analyses were performed in the *R* environment (versions 4.0.0, 4.0.3, and 4.2.1). Model reconstructions were performed with the *gapseq* software (versions 1.1 and 1.2) ^32^ Optimisations were generally conducted using the software *sybil* (versions 2.1.5 and 2.2.0) ^41^. Specifically, multi-model supplementation was conducted using the software *MicrobiomeGS2* (version 0.1.5, www.github.com/Waschina/MicrobiomeGS2) and the in-silico co-culturing was simulated on *BacArena* (version 1.8.2, commit fdb02bf7) ^36^. Both incorporate *sybil* in their pipelines. The linear programming solver was *CPLEX* (version 12.10.0, 22.1.0), based on the *R* package *cplexAPI* (version 1.4.0). For data management, the software *dplyr* (version 1.0.2, 1.0.4, and 1.0.9), *tidyr* (version 1.1.3), *tidyverse* (version 1.3.2) ^46^, and *data*.*table* (version 1.14.2) were used. For parallel computing, the software *foreach* (version 1.5.0 and 1.5.1) and *doMC* (version 12.10.0) were used. For plotting, the software *ggplot2* (versions 3.3.3 and 3.3.6) ^47^, *egg* (version 0.4.5), and *ggVennDiagram* (version 1.2.2.) were utilized.

#### Ethics statement

The human cohort data that was used in this study originate from the publication of Pryor and colleagues ^30^, whose access was provided by the *PopGen* biobank (Schleswig-Holstein, Germany) ^48^. The raw data is available from the biobank through a structured application procedure (https://www.uksh.de/p2n/Information+for+Researchers.html).

### b. Experimental Methods

#### Bacterial strains

To determine bacterial growth in mono- and co-culture either in the presence or absence of supplementation, similarly to the computational section, the two naturally associated *Caenorhabditis elegans* microbiota isolates *Pseudomonas lurida* MYb11 and *Ochrobactrum vermis* MYb71 were used. Fluorescently labeled MYb11::dTomato ^21^ and MYb71::GFP were used to distinguish the two bacterial species in a mixed community. For comparison, reverse labels were also tested, using MYb71::dTomato and MYb11::sfGFP, to ensure no impact of the fluorescent label (Supplementary Figure S2). Integrated fluorescent strains of MYb71 and the MYb11::sfGFP strain were constructed by transposon insertion at the *attTn7* site following published work ^49^. Briefly, two populations of *E. coli* SM10 strains were prepared: (i) a donor strain containing either pTn7xKS-sfGFP (pTW415) or pTn7xKS-dTomato (pTW416) plasmids, and (ii) a helper strain containing the pTNS2 (pTW10) plasmid. Liquid cultures (5ml LB) of donor, helper (both 37 °C) and target strains (26 °C) were grown with agitation overnight, subcultured and collected at an OD 0.4-0.6. Tripartite mating mixtures consisted of equal parts (1:1:1), incubated overnight at 26 °C, followed by selection on LB agar plates containing gentamicin (10 ug/mL) and IPTG (1mM). Fluorescent colonies of MYb71 were verified by PCR and sequencing using 2 pairs of MYb71-specific primers to the glmS gene region: (i) oDB097 (5’-CGCTCTGATCGATGAGACC-3’) and oDB098 (5’-TTGCGCGGCTGRTCGAC-3’), and (ii) oDB097 and a transposon WP11 primer (5’-CACGCCCCTCTTTAATACGA-3’)^49^. MYb11::sfGFP was similarly confirmed by PCR and sequencing, following our previous protocol ^21^.

Bacteria were thawed from frozen stocks for each experiment, streaked to tryptic soy agar (TSA) plates and grown for 16 h to 42 h at 25 °C. Bacterial cultures of MYb11 and MYb71 were grown in liquid nematode growth medium (liquid NGM) overnight at 28 °C in a shaking incubator. Cultures were adjusted to OD_600_ = 0.5 with phosphate-buffered saline (PBS), and co-cultures were obtained by mixing equal volumes of the adjusted cultures.

#### Supplementation in vitro

L-serine (1714.1), D-mannitol (8883.1), and L-threonine (T206.1) were purchased from Carl Roth (Karlsruhe, Germany), and GABA (A2129) from Merck (Darmstadt, Germany) and dissolved to 1 M or 0.5 M stocks in water, depending on the maximum soluble concentration. Liquid NGM was supplemented with a final concentration of 10 mM of the respective supplement. For supplementation assays *in vitro*, 5 μl of either mono- or co-culture of the bacteria was added to 95 μl of liquid NGM in a 96-well plate, in the presence or absence of supplement. Treatments were randomized with respect to plate layout. Plates were incubated at 28 °C for 24 h in an Epoch2 BioTek plate reader with double orbital shaking at 807 CPM. To quantify the colony forming units (CFU) 7 μl of different dilutions of each treatment bacterial suspension of respective mono- or co-cultures were plated on TSA plates. The plates were stored at 25 °C and colonies scored after 24 h to 48 h. Statistical calculations and plots were performed with R studio software (version 4.1.3). For all CFU assays *in vitro*, statistical comparisons were performed with Wilcoxon signed rank tests ^44^.

#### Supplementation in vivo

The *C. elegans* strain N2 was used for all *in vivo* experiments and was originally obtained from the Caenorhabditis Genetics Center (CGC). Worms were maintained on NGM plates inoculated with *Escherichia coli* OP50 at 20 °C following standard procedures ^50^. Assay plates were prepared by adding L-serine at a concentration of 0 mM, 10 mM, or 100 mM to NGM. Overnight cultures of MYb11 and MYb71 were adjusted to OD_600_ = 2.0 in PBS. 30-40 synchronized first instar larvae (L1) were grown on supplemented NGM plates inoculated with 125 μl of respective mono- or co-culture of MYb11 and MYb71. After 72 hours, bacterial lawns were sampled by pressing the circular end of a 200 μl pipette tip into the lawn, avoiding the edges and any worms. The resulting disc was homogenized in PBS with zirconia beads in a Bead Ruptor 96 (Omni International) for 3 minutes at 30 Hz.

To assess the number of CFU, worms were washed off plates using M9-buffer + 0.025% Triton X-100, washed 5 times and paralyzed with 5 mM tetramisole to stop pumping of the worms as described in ^22^. Worms were surface-sterilized following ^22^, then washed 2 times in PBS to remove residual bleach. Worms were transferred to a new tube to determine the exact worm number (∼20) and PBS was added to a final volume of 400 ml. Worms were allowed to settle and 100 μl supernatant was collected (supernatant control). Worms were homogenized in the remaining PBS using 1-mm zirconia beads in a Bead Ruptor 96 for 3 minutes at 30 Hz. Homogenized worms, lawns, and supernatants were serially diluted in PBS and plated on TSA plates. After up to 48 h at 25 °C, the colonies were scored at the appropriate dilutions and the CFUs per worm were calculated. Each colonization experiment was conducted in three independent runs and equal biological replicates. Data were pooled across runs, and a mixed model controlling for runs was used to assess the effect of L-serine supplementation on bacterial colonization in *C. elegans*. For all L-serine *in vitro* and *in vivo* experiments, statistical comparisons were performed with generalized linear models, if not otherwise mentioned. We calculated the expected proportion of MYb11 in worms as the product of MYb11 proportions available on the lawns and the observed shift in bacterial proportions due to host filtering:

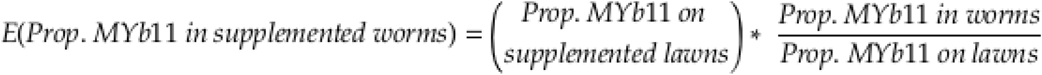

Plots and statistical analysis were produced with R studio (version 4.1.3) and edited with Inkscape (version 1.1.2).

## 5. Results

### Overlapping metabolic niches between microbiome member species prevent the use of unique uptake compounds as precision prebiotics

The simplest approach to specifically boost the abundance of a particular bacterial species in a microbial community is to supply the community with a compound that is only taken up by the target species but no other species. Hence, these compounds correspond to unique metabolic niches for the respective bacterial species. We initially tested computationally whether such unique uptake compounds typically exist in bacterial communities, using bacterial communities from a human cohort as well as *C. elegans* ^30^. We evaluated for each bacterial species of a particular community whether it was able to take up a metabolite that no other species of the community could take up and determined the fraction of bacterial species in the community that had such unique uptake compounds (see Methods). For the human cohort, the bacterial communities comprised on average 26-34 bacterial species (25% to 75% quantiles). Within this community size, on average, 5 to 20% of bacterial species were inferred to possess unique uptake compounds that could potentially be used to target those species. For smaller communities, up to 50% of bacterial species and for larger communities, only 5% to 10% had unique uptake compounds (Figure 1A). Similarly, in the *C. elegans* model system, we observed that the possibility of finding a unique compound quickly dropped with the size of the bacterial community. Thus, within a sample of 78 bacterial species isolated from *C. elegans*, we usually only identified 3-5 % of species with unique uptake compounds in communities comprising 30 or more bacterial species, which is the typical size of bacterial communities observed in *C. elegans*-derived microbiome samples ^51^ (Fig. 1B). Similarly, in the artificial CeMbio *C. elegans* model microbiome community unique uptake compounds could usually only be detected for subsets of bacterial species up to a size of six (Fig. 1C).

**Figure 1:**
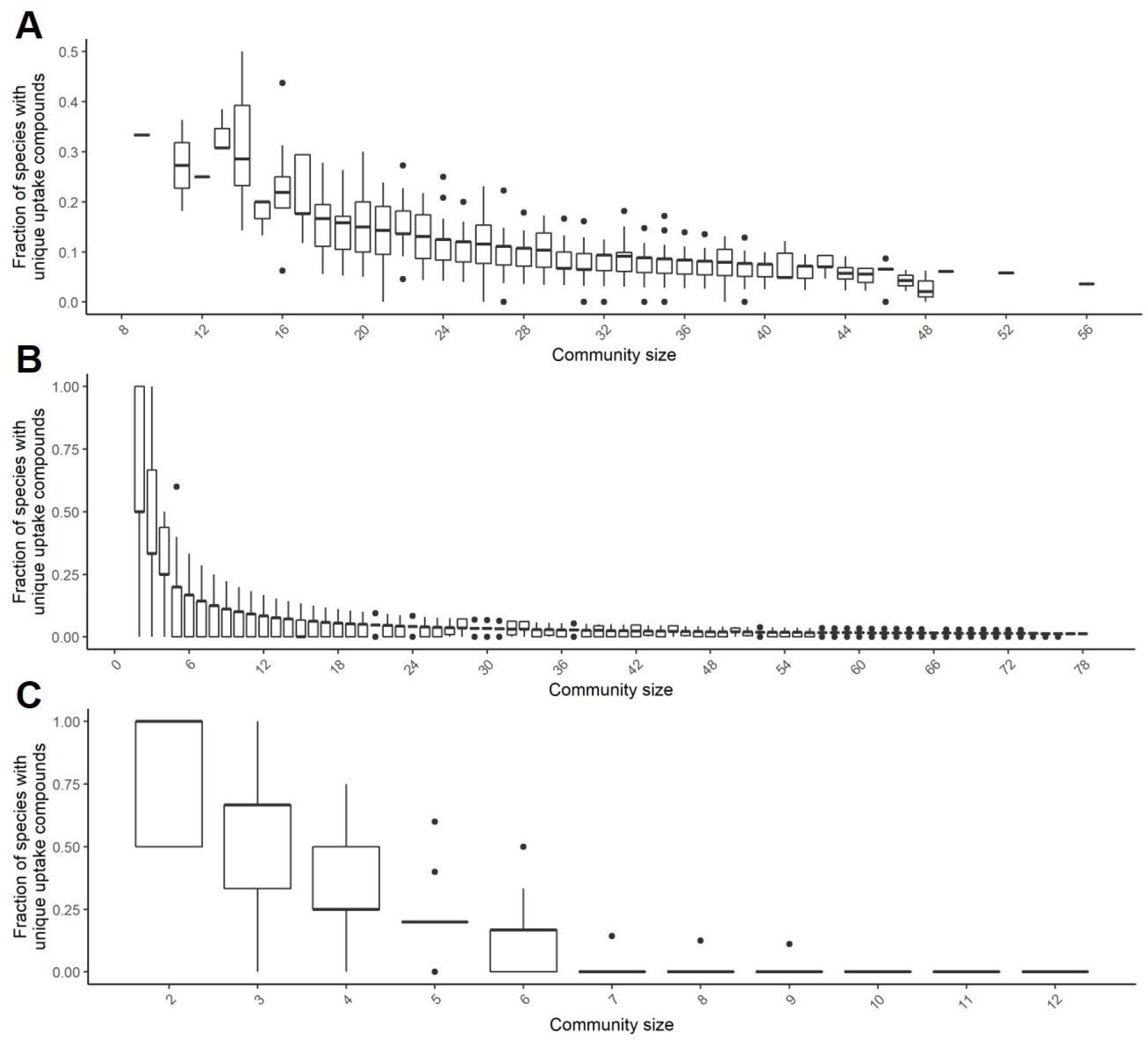
Frequency of species with unique uptake compounds in microbiomes. Fraction of species with unique uptake compounds in **(A)** human gut bacterial communities and in **(B**,**C)** *C. elegans* bacterial communities that are subsets of previously reported *C. elegans* microbiome member species.

Thus, in a typical microbiome like those analyzed above, we would expect to be able to identify unique uptake compounds only for a minority of bacterial species and compounds to target specific bacterial species are typically expected to be consumed by several species. Thus, for our identification of precision prebiotics for MYb11, we firstly excluded twenty compounds from our search that could specifically be taken up only by MYb11 in the search for a MYb-targeted prebiotic (20 compounds, see Supplementary File (Sheet: S2). Of note, MYb11 and MYb71 share the vast majority of all uptake compounds (85%), while only 10% and 5% of all uptake compounds are unique to MYb11 and MYb71, respectively (Figure 2A).

**Figure 2:**
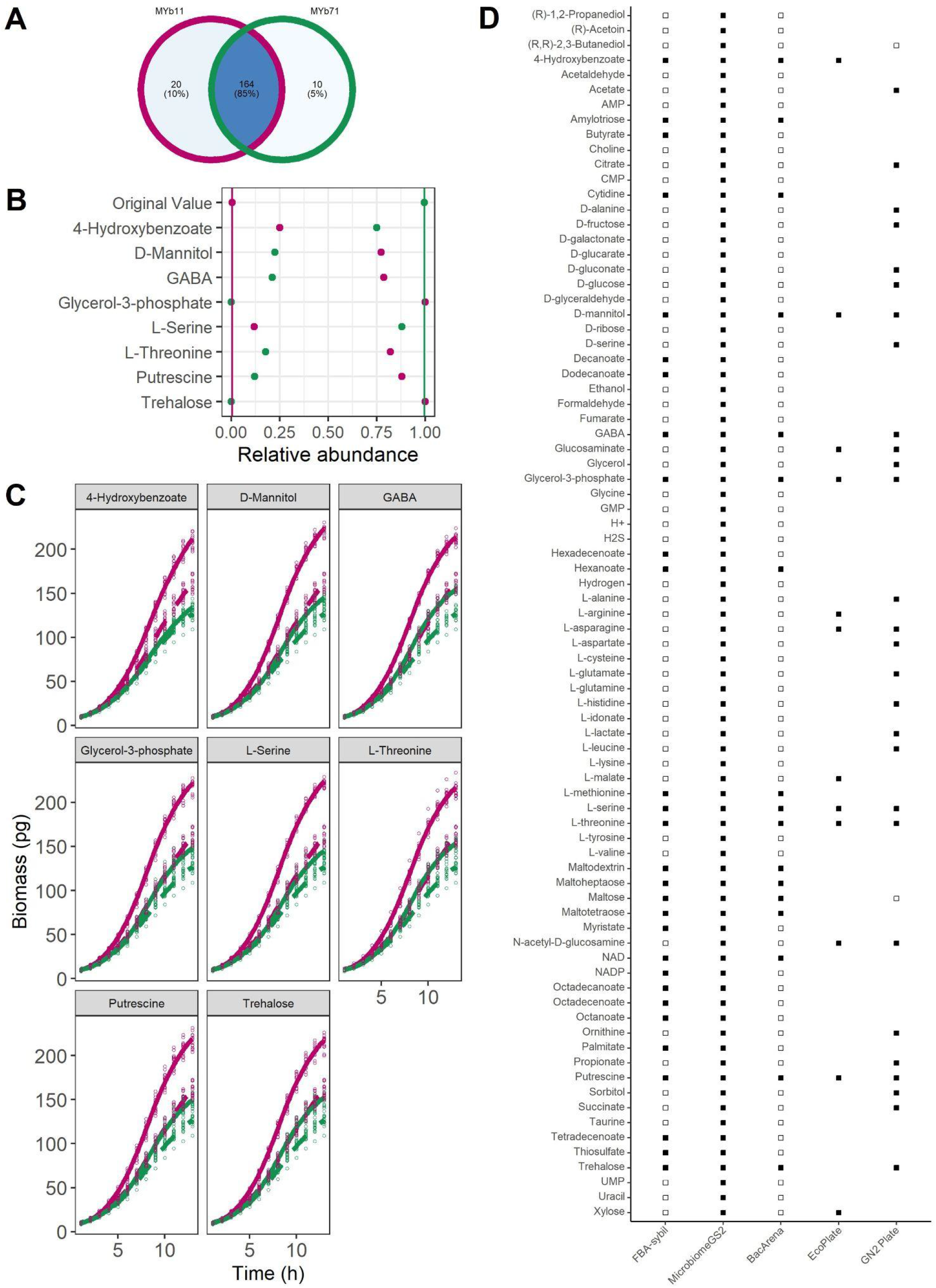
Metabolic-modeling guided identification of precision prebiotics. **(A)** Overlap of all exchange reactions across the models MYb11 and MYb71. **(B)** Community FBA supplementation of *Pseudomonas lurida MYb11* (pink) and *Ochrobactrum vermis MYb71* (green) in *MicrobiomeGS2*. Predicted relative abundance of MYb11 and MYb71 under the supplementation with 4-hydroxybenzoate, D-mannitol, GABA, glycerol-3-phosphate, L-serine, L-threonine, putrescine and threhalose shown in dots, lines represent predicted relative abundance without any supplementation. **(C)** *In-silico* supplementation of *Pseudomonas lurida MYb11* (pink) and *Ochrobactrum vermis MYb71* (green) in *BacArena*. Predicted growth curves of MYb11 and MYb71 either with 10 mM of 4-hydroxybenzoate, D-mannitol, GABA, glycerol-3-phosphate, L-serine, L-threonine, putrescine or trehalose(solid line) or without supplementation (dashed line). **(D)** Comparison of proposed compounds with different computational approaches and previously published information, using *BacArena*, community FBA/*MicrobiomeGS2*, individual FBA, and growth data in *Biolog* plates ^22,27^. Empty squares represent cases where no growth was observed in Biolog data. Missing squares indicate compounds not present on Biolog plates.

### Computational identification of precision prebiotics targeting Pseudomonas lurida MYb11

In order to derive precision prebiotics for *Pseudomonas lurida* MYb11, we used its metabolic network as input for flux balance analysis and two distinct community modeling approaches apart from the basic approach of FBA. The first one is based on *BacArena* that incorporates an individual-based modeling approach and the other one is so-called community FBA implemented in *MicrobiomeGS2*. These two modeling approaches differ in their underlying assumptions. In *BacArena* the two bacteria are modeled by their individual metabolic networks and fluxes are predicted by maximizing the growth rate of individual bacteria. In community FBA, in turn, the metabolic networks of the individual bacterial species are merged into a community model for which the sum of growth rates of both bacterial species are maximized. Thus, when using *BacArena*, metabolic interactions between species typically arise from one species secreting an end-product that another species can metabolize. In community FBA, in turn, the optimization of total growth assumes that bacterial species coordinate their metabolic fluxes so that overall growth is maximized.

Using FBA, we identified 29 compounds that could increase the growth of MYb11 in single growth. Employing the community modeling approaches, we identified 81 compounds that increased the growth of MYb11 over MYb71 in community FBA and 17 compounds using *BacArena*. In order to determine whether specific compound classes were enriched among the MYb11-supporting compounds, we performed an enrichment based on HMDB annotation ^45^ across the three different simulation types. We found that growth-supporting compounds for MYb11 were enriched for amino acids (Fisher’s exact test *p*-value=2.2×10^−2^) and fatty acids (Fisher’s exact test *p*-value=3.16×10^−5^). Interestingly, the growth dynamics of the models when they were part of a community model (Figure 2B) differ from the growth dynamics of the models when they were treated as individual organisms (Figure 2C). For instance, MYb71’s growth rate was much higher in the community model, i.e., when coupled to the model MYb11. This is a typical observation when using community FBA when the growth rate of the entire community is maximized. Especially for small communities, this often leads to one community member that most efficiently turns the growth medium into biomass to grow strongest with only little growth for other community members. On the other hand, in the *BacArena* simulations, we observed that most of those compounds boosted the growth of both species in single growth, however, the effect was larger for MYb11 than MYb71 (Figure 2C). Of the identified compounds, 17 compounds were shared across the three simulations. For eight of these, uptake by MYb11 was also supported by *Biolog* plates ^22,27^ (Figure 2D). Based on the available literature information on the role of these compounds in longevity of the host ^52,53^ or physiology ^54^, as nutritional compounds ^22,55^, or compounds that are involved in host-microbial interactions ^54,56^, we selected four candidates for further experimental testing as precision prebiotics to specifically increase the abundance of MYb11 based.

### In vitro validation of predicted precision prebiotics

We experimentally tested the ability of four of the predicted precision prebiotics compounds, L-serine, L-threonine, GABA, and D-mannitol, to specifically increase the growth of MYb11 in co-culture with MYb71. L-serine is a dietary amino acid that has been involved in nutrition-drug-host-microbial interactions ^56^ and can support the lifespan of the host ^53^, L-threonine, an essential amino acid ^55^, through its catabolite methylglyoxal (in low amounts) can also have a positive effect on host’s lifespan ^52^. GABA, apart from being a bacterial product, is protective for the host’s neurons ^54^. Lastly, D-mannitol is a commonly-used sugar alcohol^22^, therefore its usage as a nutritional compound would make its usage as a supplement a safe option for the host.

We supplemented MYb11 and MYb71 in mono- as well as co-culture and measured bacterial growth *in vitro* in liquid NGM after 5 h (Supplementary Figure S1) and 24 h. Independently of the added supplement or the mono- or co-culturing, MYb11 and MYb71 showed increased growth in mono- and co-cultures when supplemented with 10 mM L-serine, L-threonine, GABA, as well as D-mannitol after 24 h (Figure 3A). Additionally, to quantify bacterial growth we used fluorescently-labeled strains. CFU of mono- and co-culture were plated and counted after 24 h growth in liquid culture in the presence (10 mM) or absence (0 mM) of the four compounds L-serine (Figure 3B), L-threonine (Figure 3C), GABA (Figure 3D) and D-mannitol (Figure 3E). MYb11 and MYb71 showed increased colony numbers in mono-culture when supplemented with L-serine and L-threonine compared to no supplementation (Figure 3B, C, Wilcoxon signed rank test, *p* = 0.001, *p* = 0.006, *p* = 0.007, *p* = 0.032, respectively). GABA supplementation led to an increase in growth of MYb11 in mono-culture, but not MYb71 (Figure 3D, Wilcoxon signed rank test, *p* < 0.001, *p* = 0.922, respectively), while D-mannitol had no effect on the growth of MYb11 nor MYb71 in mono-culture (Figure 3E, Wilcoxon signed rank test, *p* = 0.259, *p* = 0.661, respectively). However, all four compounds, L-serine, L-threonine, GABA and D-mannitol specifically promoted MYb11 growth in co-culture over MYb71 *in vitro* and thus confirmed the computational predictions (Figure 3C-E, Wilcoxon signed rank test, *p* = 6.87e-06, *p* < 0.001, *p* = 7.15e-06, *p* = 7.07e-06, respectively)

**Figure 3:**
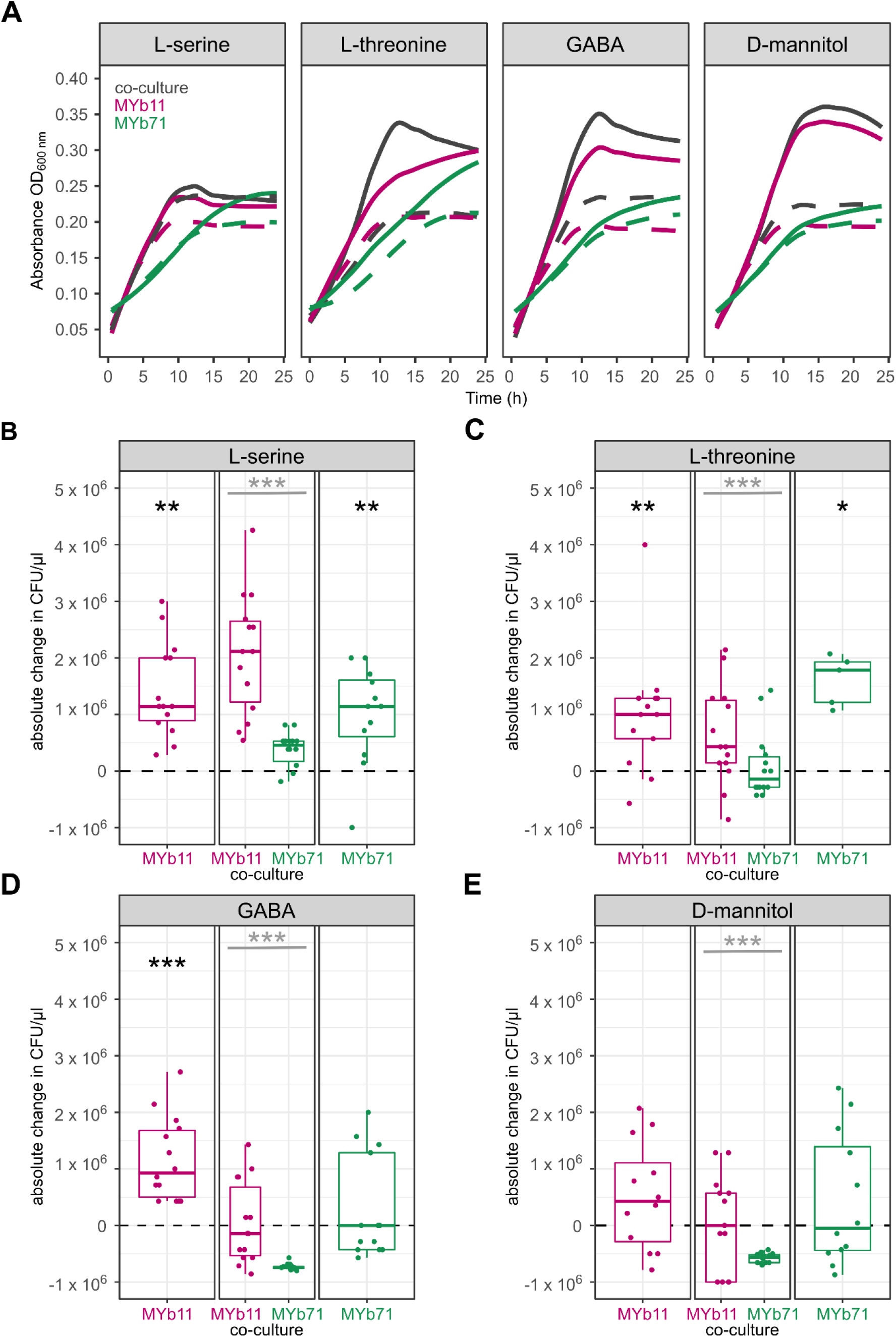
*In vitro* growth of *Pseudomonas lurida* MYb11 and *Ochrobactrum vermis* MYb71 in mono- and co-culture with four different supplements for 24 h. **(A)** Growth curves of co-cultures (gray) or mono-cultures of MYb11 (pink) and MYb71 (green) in liquid NGM for 24 h either with 10 mM of L-serine, L-threonine, GABA, or D-mannitol (solid line) or without supplementation (dashed line). **(B, C, D, E)** Colony-forming units (CFU/μl) in mono-cultures of MYb11 (left) and MYb71 (right) or in co-culture (middle) after 24 h of 10 mM of the respective supplement. Shown are boxplots with the median as a thick horizontal line, the interquartile range as box, the outer quartiles as whiskers, and each replicate depicted by a dot. Every replicate was normalized by subtracting the non-supplemented median of the respective bacteria (dashed line). Black asterisks indicate statistical comparisons between supplemented and non-supplemented median, gray asterisks indicate statistical comparisons between supplemented medians of MYb11 and MYb71. n = 5-14.

### Concentration-dependent effect of L-serine supplementation in vitro

The statistical analyses suggested that L-serine is the most promising precision prebiotic for further analysis among the compounds validated *in vitro*. We repeated the *in vitro* supplementation experiment across a concentration gradient to determine the optimal dose for enrichment of MYb11. Overall, the MYb11 growth as well as MYb71 growth in monoculture at 10 mM and 100 mM over time were increased following the increase in L-serine concentrations (Figure 4A). When quantifying the growth with supplementation at the same concentration gradient (Figure 4B), then starting with 1 mM did not significantly impact the abundance of MYb11 proportions (GLM, *p* = 0.778), nor MYb71 (GLM, *p* = 0.69). However, supplementation with 10 mM led to an enrichment of MYb11 (GLM, *p* < 0.001), thereby indicating a significant difference in proportions in co-culture between MYb11 and MYb71 (GLM, *p* = 5.77E-07). Essentially identical results were obtained at 100 mM, including an MYb11 enrichment (GLM, *p* = 0.004), and a significant difference between MYb11 and MYb71 proportions (GLM, *p* = 7.93 E-05).

**Figure 4.**
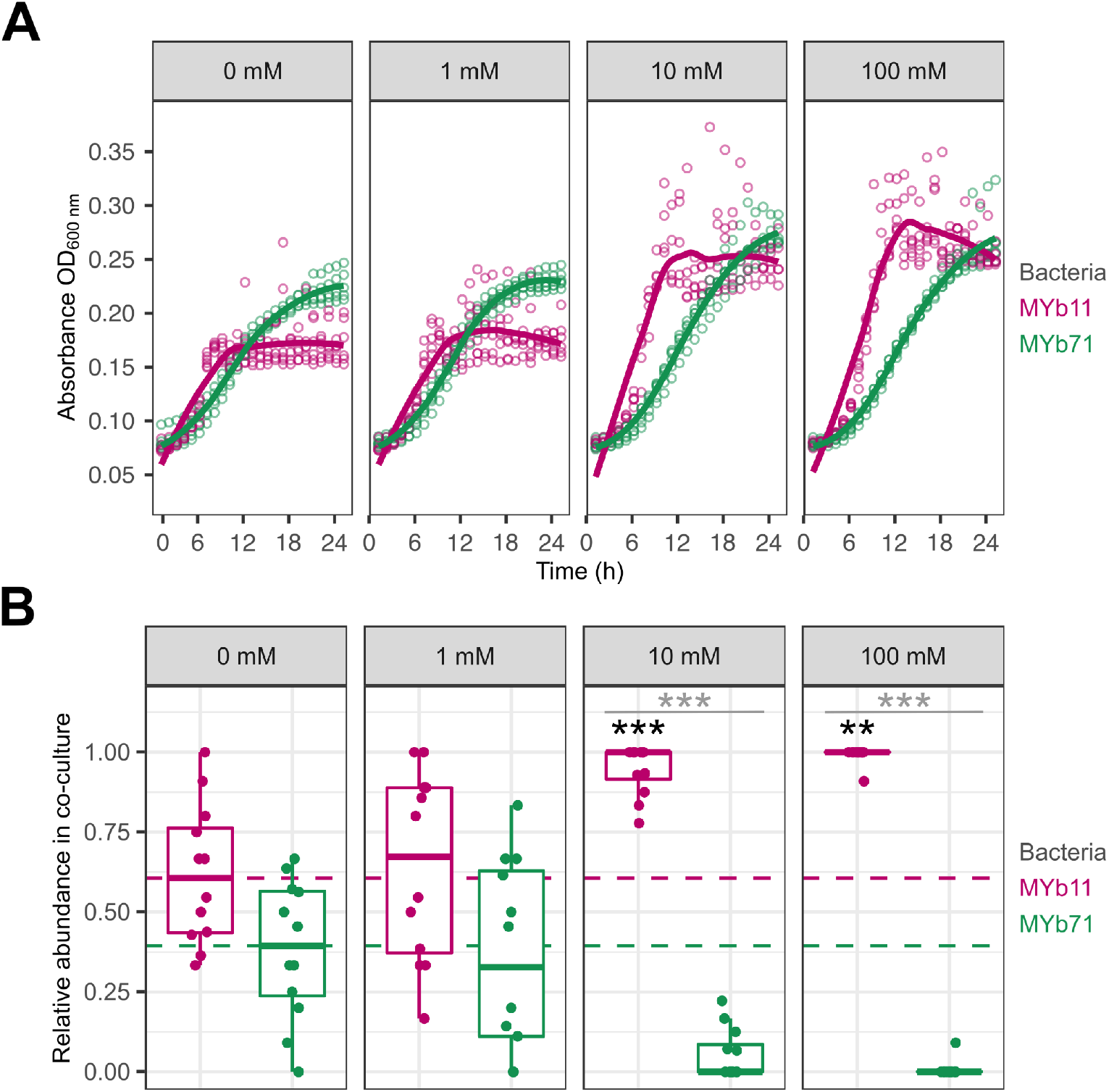
Concentration-dependent effect of L-serine supplementation on bacterial growth *in vitro*. **(A)** Bacterial growth was measured based on optical density (OD_600_) of each species in mono-culture in liquid nematode growth medium (NGM) with the respective L-serine supplementation of 0 mM, 1 mM, 10 mM or 100 mM. Points represent individual replicates, while lines indicate the mean, n = 12. **(B)** Relative abundance of MYb11 (pink) and MYb71 (green) in co-cultures grown for 24 h with the respective supplement concentration in liquid NGM, based on colony counts of fluorescently labeled bacteria. Shown are boxplots with the median as a thick horizontal line, the interquartile range as box, the outer quartiles as whiskers, and each replicate depicted by a dot. Every replicate was normalized by subtracting the non-supplemented median 0 mM (dashed line) of the respective bacteria. Statistical differences were determined by GLM and are indicated by asterisks (*** *p* < 0.001). Black asterisks indicate statistical comparisons between supplemented and non-supplemented median, gray asterisks indicate statistical comparisons between supplemented medians of MYb11 and MYb71. n = 9-12.

### Serine supplementation alters bacterial colonization of C. elegans

Finally, we tested whether L-serine supplementation to solid growth medium would alter bacterial proportions in the host *C. elegans*. Synchronized populations of L1 larvae were pipetted onto bacterial lawns of MYb11 and MYb71 on solid NGM plates supplemented with either 0 mM, 10 mM, or 100 mM L-serine. After 72 h, adult worms were collected to quantify bacterial proportions. Supplementation at 10 mM L-serine did not change the proportion of MYb11 in co-culture in worms compared to non-supplemented media (GLM, *p* = 0.459, Figure 5A left panel). However, L-serine supplementation at 100 mM increased the proportion of MYb11 in the bacterial community in worms compared to non-supplemented media (GLM, *p* < 0.001,Figure 5A right panel). We next asked whether the effect of supplementation on bacterial proportions in the host simply reflected changes in the proportion of bacteria that were available to them on the lawn (i.e., their diet). The relative abundance of MYb11 on lawns did not change by 10 mM supplementation (GLM, *p* = 0.61, Figure 5A, left panel) compared to non-supplementation. However, the relative abundance of MYb11 was approximately 60% in non-supplemented media and increased to approximately 75% with 100 mM L-serine supplementation (GLM, *p* < 0.002, Figure 5B, right panel). The observed proportions of MYb11 in worms at 100 mM serine were significantly higher than expected based on proportions in the lawn and thus colonization probability compared to MYb71 (Chi-squared test, χ^2^ = 10.244, df = 1, *p* = 0.0013; see Methods). This suggests that the provision of serine enriches MYb11 in the host independently from the enrichment observed on the lawn and that serine allows MYb11 to overcome its growth-disadvantages in competition with MYb71 in the host.

**Figure 5.**
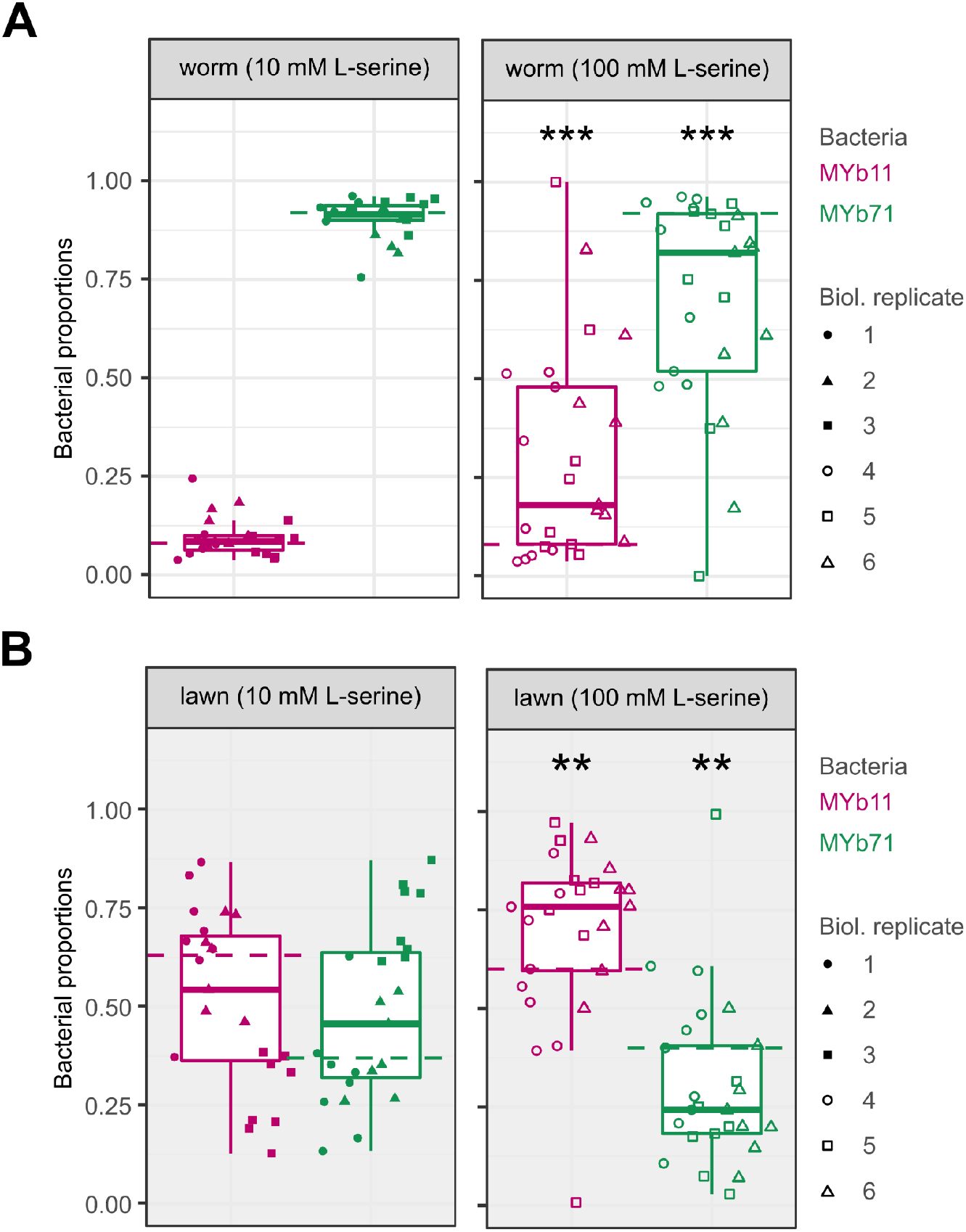
L-serine supplementation alters bacterial abundance in *C. elegans*. **(A)** Adult worms were collected from solid NGM plates, which had been inoculated with co-culture and supplemented with 10 mM L-serine (left panel) or 100 mM (right panel) L-serine. Worms were washed and surface-sterilized to remove residual bacteria, then homogenized in a buffer. Bacterial proportions were quantified based on colony counts. **(B)** Bacterial lawns were sampled by cutting an agar disc of standardized size from NGM plates supplemented with 10 mM (left panel) or 100 mM L-serine (right panel). Discs were homogenized in buffer, and CFU were quantified. **(A, B)** Shown are boxplots with the median as a thick horizontal line, the interquartile range as box, the outer quartiles as whiskers, and each biological replicate depicted by a symbol. Every replicate was normalized by subtracting the non-supplemented median 0 mM (dashed line) of the respective bacteria. Statistical differences were determined by GLM and are indicated by asterisks (*** *p* < 0.001, ** *p* < 0.01) and show comparisons between supplemented and non-supplemented median

## 6. Discussion

Although many factors are known to influence microbiome composition (e.g., dietary patterns, lifestyle or medication ^57^), the mechanisms causing these changes is often not fully understood and often a large number of bacterial species are affected. We propose that the most effective way to modulate the composition of the microbiome is by modulating or enriching the abundance of an already existing bacterial species with health-beneficial effects using precision prebiotics. A direct approach to identify such precision prebiotics would be to provide compounds that are uniquely taken up by the bacterial target species. However, through an analysis of *C. elegans* and human microbiome data, we were able to show that such unique uptake compounds typically only exist for few species in a microbial community and especially for larger communities there are no such compounds at all. Thus, in a typical microbiome, precision prebiotics have to be identified from compounds that are potentially taken up by several species. To identify such compounds, we used genome-scale metabolic networks of two bacterial members from the *C. elegans* microbiome, *Ochrobactrum vermis* MYb71, and the host-protective *Pseudomonas lurida* MYb11, in conjunction with constraint-based modeling to identify compounds that specifically increased the growth of MYb11. Using HMDB annotation ^45^, we found that amino acids and fatty acids were enriched as predicted growth-supporting compounds for MYb11 which is in agreement with previous observations of the preference of bacteria of the genus *Pseudomonas* for these compounds ^58–61^. Considering compounds that were identified across all three simulation approaches and cross-referencing with *Biolog* data revealed that 4-hydroxybenzoate, D-mannitol, γ-aminobutyric acid (GABA), glycerol-3-phosphate, L-serine, trehalose, L-threonine, and putrescine are potential precision prebiotics for MYb11. We experimentally tested four of these predicted compounds, L-serine, L-threonine, D-mannitol, and GABA *in vitro* and confirmed their positive effect specifically on the growth of MYb11. Moving into the *C. elegans* host system, we were able to show that L-serine is also able to selectively boost the abundance of MYb11 *in vivo*, when the host was colonized with both bacterial species. Interestingly, while 10 mM of L-serine were sufficient for an enrichment of MYb11 in batch culture, 100 mM were necessary in the growth experiments with the host. This suggests differences in either bacterial uptake kinetics for L-serine or bacterial growth in general between batch growth and growth on agar plates. Moreover, while MYb11 was strongly enriched in worms to similar levels like MYb71 even though it had a very low abundance in the lawn, this enrichment was even stronger when supplying L-serine. Thus, our proof-of-principle work demonstrates that constraint-based microbial community modeling approaches are an effective tool for the future development of targeted microbiome therapies using precision prebiotics.

The effect of serine and threonine on the host and/or on its microbiome has been also documented in the literature. For instance, threonine is an essential amino acid for the host ^55^. Longevity effects on the host have been shown also after supplementation with threonine ^62^ and serine ^53^, while dietary serine reduced host fitness through acting on bacterial metabolism, when it was supplemented along with the chemotherapeutic 5-fluoro 2′deoxyuridine ^56^. Moreover, serine promotes the growth of *E. coli* LF82 in the mouse gut ^63^, which shows the potential of this compound to alter the abundance of multiple species. In accordance, we also observed effects of serine on MYb71 growth, which was, however, stronger for MYb11 if they were grown in community.

One important challenge to specifically target the microbiome with nutritional supplements is to prevent host uptake or metabolization of the supplements ^64^. Different approaches to achieve restricted bioavailability, such as the supplementation of non-digestible carbohydrates ^65^ or coating of supplements with compounds that dissolve during passage through the digestive tract have been developed ^66–68^. Moreover, as indicated by our analysis of concentration-dependent effects of supplementation in the case of serine, it might be necessary to control the specific concentrations of a compound that are being released and their stability over time in the gut. To maintain relatively constant concentrations or to deviate from physiological constraints, advanced formulations (such as nanotechnology ^69^, delayed release and other drug delivery technologies ^70,71^) could be employed.

Although our combined computational-experimental approach can serve as a blueprint for future studies aimed at the identification of precision prebiotics, several factors should still be taken into consideration. As we showed, it is nearly impossible to identify a compound that is uniquely taken up by only one species within a community. This point becomes apparent, if we consider the numerous different species that reside in various hosts. For instance, the synthetic bacterial community of *C. elegans, CeMbio* ^22^ has 12 members, while the human-related bacterial community AGORA has more than 800 species ^34^. Thus, an important next step is to scale up our approach to more complex communities. Furthermore, since the physiology of worms cannot be directly compared to the physiology of humans, the efficacy and effectiveness of the approach (see more on the topic ^72^) need to be studied in more detail. At this point, the complexity of the regulatory affairs in various jurisdictions (e.g., the definition of nutritional supplementation vs. medication, safety concerns) should also not be underestimated ^73^. Taking everything into consideration, we showed that the design of targeted supplementation strategies to modulate the microbiome is not only feasible but also promising in terms of future microbiome-based treatment.

## 7. Authors’ Contributions

Conceptualization: CK, GM, NO, HS

Methodology: DB, KD, RD, IKH, CK, GM, NO, CP, BSS, HS, JT, JZ

Software: RD, IKH, CK, GM, CP, JT, JZ

Formal analysis: RD, IKH, CK, GM, NO

Investigation: RD, IKH, CK, GM, NO, CP

Resources: DB, KD, AF, CK, ML, BSS, HS

Data Curation: RD, IKH, GM

Writing - Original Draft: IKH, CK, GM

Writing - Review & Editing: All co-authors

Visualization: RD, IKH, GM

Supervision: CK, NO, HS

Project administration: CK, GM

Funding acquisition: KD, RD, CK, GM, BSS, HS

## 8. Acknowledgements

We acknowledge support by the German Research Foundation within the collaborative research center “Origin and Function of Metaorganisms”, CRC1182, sub-project A1 to CK, KD, and HS; the research unit miTarget (FOR5042) to CK and the excellence cluster “Precision medicine in chronic inflammation” (EXC2167) to CK and HS. CK moreover acknowledges support by the German Ministry for Education and Research (E:Med iTREAT, support code 01ZX1902A). GM received support through a Young Investigator Award by the CRC1182. RD received support through a Short-Term Research Grant from the Deutscher Akademischer Austauschdienst (German Academic Exchange Service). We also acknowledge funding via NIH grant DP2DK116645, NASA grant 80NSSC22K0250, and JGI/DOE grant CSP-503338 (all to BSS). This research was supported in part through high-performance computing resources available at the Kiel University computing centre. *C. elegans* N2 was originally provided by the Caenorhabditis Genetics Center (CGC; https://cgc.umn.edu/), which is funded by the NIH Office of Research Infrastructure Programs (P40 OD010440).

## 10. Data availability

The R scripts and the raw data to conduct the simulations are deposited in GitHub (www.github.com/maringos/Nutritional-Supplements-Prediction) (version / commit: 36ff648). The growth rate changes of the supplementation experiments with FBA *Sybil* and MicrobiomeGS2 can be found in the Supplementary File (Sheets: S3, S4). The bacterial biomass data of the *BacArena* simulations can be found in the Supplementary File (Sheet: S5).

## 11. Supplement

**Figure S1:**
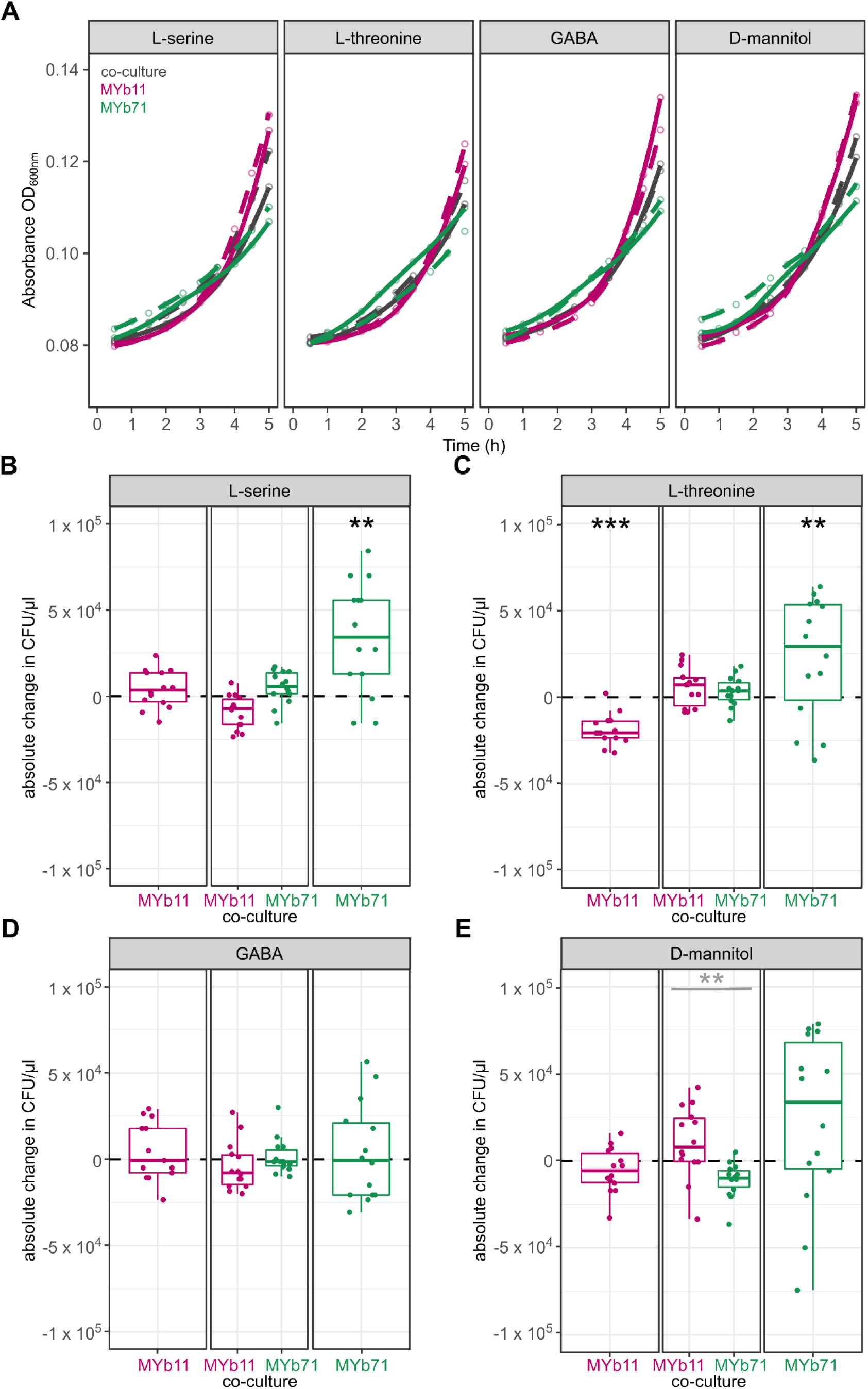
*In vitro* growth of *Pseudomonas lurida* MYb11 and *Ochrobactrum vermis* MYb71 in mono- and co-culture in the presence and absence of four different supplements for 5 h. **(A)** Growth curves of mono- and co-cultures (gray) of MYb11 (pink) and MYb71 (green) in liquid NGM for 5 h either with 10 mM of L-serine, L-threonine, GABA, and D-mannitol (solid line) or without supplementation (dashed line). **(B, C**, **D, E)** Colony-forming units (CFU/μl) in mono-cultures of MYb11 (left) and MYb71(right) or in co-culture (middle) after 5 h of 10 mM of the respective supplement. Shown are boxplots with the median as a thick horizontal line, the interquartile range as box, the whiskers as vertical lines, and each replicate depicted by a dot. Every replicate was normalized by subtracting the non-supplemented median (dashed line) of the respective bacteria. Statistical differences were determined by Wilcoxon signed rank test and are indicated by asterisks (*** *p* < 0.001, ** *p* < 0.005, * *p* < 0.05). Black asterisks indicate statistical comparisons between supplemented and non-supplemented median, gray asterisks indicate statistical comparisons between supplemented medians of MYb11 and MYb71. n = 5-14.

**Figure S2.**
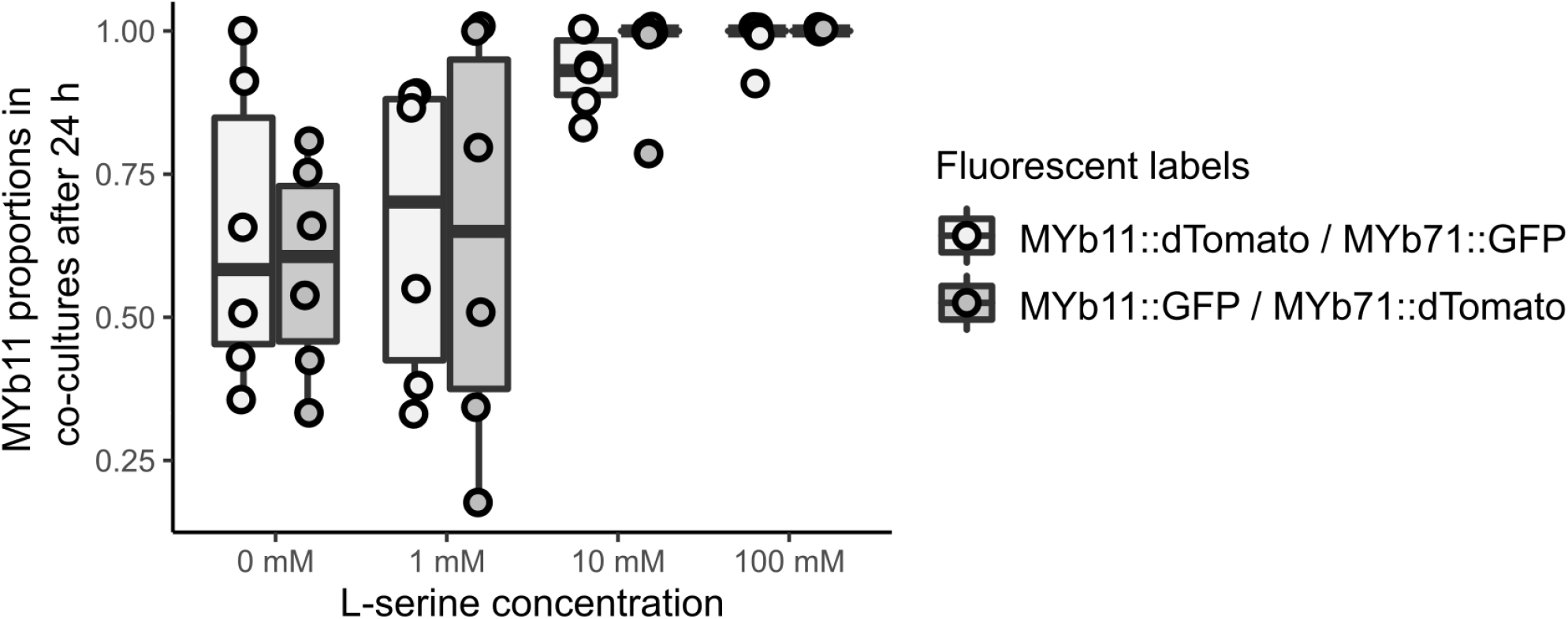
No difference in relative growth rates of differentially labeled bacterial strains. Colony forming units of co-cultures of either MYb11::dTomato/MYb71::sfGFP or MYb11::GFP/MYb71::dTomato were quantified after 24 h of growth in liquid NGM. There was no difference between the two fluorescent labeling systems at any concentration of L-serine (GLM, *p* = 0.528 for main effect of fluorescence; and *p* = 0.379 for interaction between fluorescence and concentration).

